# Using Autoregressive-Transformer Model for Protein-Ligand Binding Site Prediction

**DOI:** 10.1101/2025.03.11.642700

**Authors:** Mahdi Pourmirzaei, Salhuldin Alqarghuli, Farzaneh Esmaili, Mohammadreza Pourmirzaei, Mohsen Rezaei, Dong Xu

## Abstract

Accurate prediction of protein-ligand binding sites is critical for understanding molecular interactions and advancing drug discovery. Existing computational approaches often suffer from limited generality, restricting their applicability to a small subset of ligands, while data scarcity further impairs performance, particularly for underrepresented ligand types. To address these challenges, we introduce a unified model that integrates a protein language model with an autoregressive transformer for protein-ligand binding site prediction. By framing the task as a language modeling problem and incorporating task-specific tokens, our method achieves broad ligand coverage while relying solely on protein sequence input. We systematically analyze ligand-specific task token embeddings, demonstrating that they capture meaningful biochemical properties through clustering and correlation analyses. Furthermore, our multi-task learning strategy enables effective knowledge transfer across ligands, significantly improving predictions for those with limited training data. Experimental evaluations on 41 ligands highlight the model’s superior generalization and applicability compared to existing methods. This work establishes a scalable generative AI framework for binding site prediction, laying the foundation for future extensions incorporating structural information and richer ligand representations. The code, model, and datasets are available at this link.

## 1 Introduction

Accurate prediction of protein-ligand binding residues is crucial for understanding biological processes such as gene regulation, signal transduction, and antigen–antibody interactions Zhao et al. (2020); Dhakal et al. (2022); Xia et al. (2024). Identifying protein-ligand interaction sites is also critical for drug discovery and design Xia et al. (2020). Despite its importance, identifying potential binding sites for specific ligands remains a significant challenge, especially in cases where the high-resolution 3D structure of a protein is unavailable Yu et al. (2013). Experimental methods like nuclear magnetic resonance and absorption spectroscopy, though reliable, are costly and time-consuming, underscoring the need for efficient computational solutions Yuan et al. (2022).

Sequence-based and 3D structure-based prediction methods are particularly valuable for addressing challenges posed by metal ions, which play vital roles in protein structural stability, metabolism, signal transport, and catalysis Xia et al. (2020); Essien et al. (2023). However, the small size and high versatility of metal ions introduce additional complexities.

Despite advancements in computational methods for protein-ligand binding residue prediction, several challenges remain. Current approaches often lack generality, restricting their applicability to a limited number of ligands Zhao et al. (2020); Xia et al. (2022); Fang et al. (2023); Wang et al. (2024). This limitation is particularly problematic given the vast diversity of ligands encountered in biological systems. Furthermore, the pervasive issue of data scarcity for certain ligands further hampers the effectiveness of conventional methods, as they struggle to generalize on most ligands with limited training samples Abdelkader & Kim (2024); Gangwal et al. (2024); Harren et al. (2024). Recent deep learning-based techniques also often overlook the potential of considering biochemical features as an interpretation measurement, which could provide deeper insights into model mechanisms in correlating different ligands, and to some extent, enhance interpretability in protein-ligand binding site prediction.

To address these gaps, this work introduces a unified model incorporating an autoregressive transformer connected to a pre-trained protein language model, ESM-2 Lin et al. (2023), for predicting protein-ligand binding sites directly from protein sequences for dozens of ligands (Figure 1). Inspired by Prot2Token Pourmirzaei et al. (2024; Pourmirzaei et al. 2025), we incorporate self-supervised pre-training tasks to provide the initial weights of the autoregressive component of the model, enabling significantly improving its performance on binding site prediction. Our approach sets an important foundation for broader and more interpretable protein-ligand binding site prediction, laying the groundwork for future methods that can extend to even more diverse ligand types and effectively address data scarcity issues.

**Figure 1.**
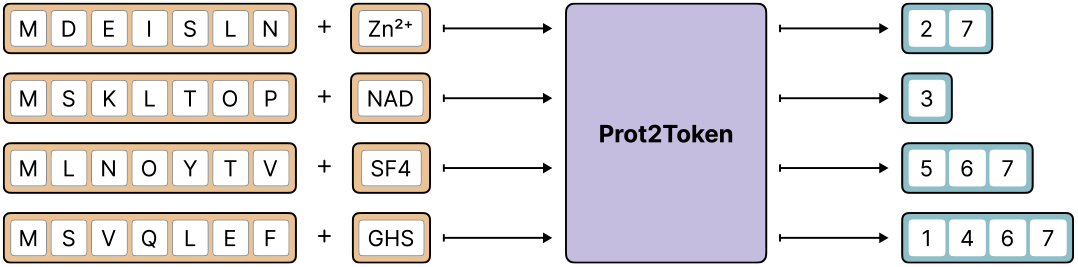
High-level overview of our approach in protein-ligand binding site prediction. The model gets protein sequences and task tokens (refer to the type of ligands) to predict the residue indices of positive sites.

Our work introduces several novel contributions which are summarized as follows:

1. By framing binding site prediction as a language modeling task and employing an autoregressive transformer model as the predictor, we achieve unprecedented generality compared to prior research, supporting 41 ligands. This approach significantly extends the applicability of protein-ligand binding site prediction methods that rely only on protein sequences for prediction.
2. By comparing biochemical features with model-generated embeddings for each ligand, we offer a comprehensive perspective on ligand relationships. Clustering and correlation analyses confirm that these embeddings capture meaningful biochemical similarities, providing deeper insights into protein-ligand interactions and improving predictions for ligands with limited sample sizes.
3. We utilize a multi-task learning approach with a unified model to enable effective knowledge transfer between ligand predictions. By leveraging embedding similarities among ligands, our method significantly enhances performance, addressing challenges posed by data scarcity and boosting predictions for underrepresented ligands.

## 2 Related Work

Computational methods for protein binding site prediction are becoming increasingly important, as the number of unknown protein 3D structures far exceeds the number of known ones Zhang et al. (2024b). Even when a protein’s 3D structure is known, the identification of all its binding ligands remains incomplete or uncertain Wang et al. (2024).

We broadly categorize protein binding site prediction methodologies into two main clusters: Standalone methods, which propose new approaches, and ensemble methods, which combine existing methods to achieve better performance. We further divide standalone methods into three subcategories: classical, deep learning, and large language methods. For a more detailed discussion of these categories and related work, refer to Appendix A.1.

## 3 Method

Our method is based on a modified version of the Prot2Token architecture, where a causal (autoregressive) Transformer, referred to as the decoder, is connected to a pre-trained bidirectional Transformer Vaswani (2017), referred to as the encoder. We initialize the encoder using the ESM-2 650M architecture and its pre-trained weights. The output of the encoder serves as context for the decoder through cross-attention. Additionally, we employ two distinct tokenizers and embedding tables for the encoder and decoder components to accommodate their unique roles in the model (Figure 3).

To guide the decoder’s predictions for specific tasks, we incorporate task tokens as part of its input. Unlike the original Prot2Token model, which utilized a pre-trained chemical language encoder, we replace this component by directly converting each ligand type into a corresponding task token provided to the decoder. This approach informs the model about the ligand type solely through the task token, eliminating reliance on the chemical language encoder.

To enhance the sequence representation of amino acids positions, we introduce a learnable positional embedding layer. The embeddings from this layer are summed with the sequence embeddings produced by the encoder. This addition compensates for the diminished positional information in the encoder’s sequence embeddings, ensuring the model effectively captures each amino acid positions during prediction.

The autoregressive transformer factorizes the joint probability of a sequence *x* = (*x*_1_, *x*_2_, …, *x*_*T*_) into a product of conditional probabilities, i.e.,

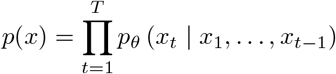

The model is trained by minimizing the negative log-likelihood of the observed tokens, expressed as

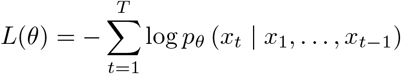

where *θ* denotes the model parameters. By using a causal mask, each token *x*_*t*_ can only attend to the tokens *x*_1_, …, *x*_*t*−1_, thus enforcing the autoregressive property and enabling the model to learn rich contextual representations of the input sequence.

We extend the standard autoregressive modeling objective by introducing token-level weights *w*_*t*_ to control the contribution of each token to the loss. In particular, we set *w*_1_ = 0 to avoid penalizing the model for predicting the first token (i.e., the prompt), and we keep *w*_*t*_ for *t* ≥ 2 adjustable so that non-prompt tokens can have varying degrees of importance. Concretely, the training objective becomes

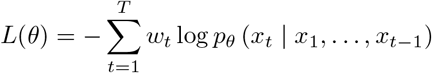

where *θ* are the model parameters and *w*_*t*_ ∈ [0, ∞) is a user-specified weight for token *x*_*t*_. This setup allows us to fine-tune the learning process by placing greater or lesser emphasis on specific tokens of each task, while completely removing the prompt token from the loss by assigning it a zero weight. More details about the architecture are provided in Appendix A.2.

### Self-supervised Pre-training of Decoder

In contrast to using a pre-trained encoder like ESM-2 weights, the decoder part is always initialized using random weights. However, the original Prot2Token paper showed that some tasks can benefit from being jointly trained with self-supervised tasks during the training of phosphorylation post-translation modification task. We speculate the main reason for this is that the decoder weights must be able to understand the structure of labels (implicit inductive biases) before making predictions. In some cases (tasks) where the vocab sizes of labels are larger (like phosphorylation), the number of samples is not sufficient for understanding those implicit biases and that could severely deteriorate the performance of the model in making predictions. In this paper, we addressed the issue by presenting a self-supervised pre-training step to provide the initialization weights of the decoder for the target task. In these self-supervised tasks, we prepared the sequences of amino acids as the input, with the labels of pinpointing the positions of specific amino acid types. For example, in sequences containing the amino acid *‘S’*, such as *“MSGLSNYT”*, we labeled the locations of *‘S’* as the target, resulting in a sequence of indices like {2,5}. Similarly, we built 20 self-supervised tasks given each amino acid as one task. The important point about these types of self-supervised tasks is that they are free to create, and consequently, no human labeling is required. By doing so, we explicitly train the decoder to recognize amino acid types and their positional indices, enabling it to construct more informative embedding tables for these vocabularies. This self-supervised learning approach is only effective when the protein encoder remains frozen; otherwise, it introduces a shortcut learning effect that can cause the model to collapse

### Tokenization

We adopted the original Prot2Token tokenization strategy by representing the positions of positive sites as a single token in ascending order. For instance, consider a protein sequence such as ‘MNSSKYKAPTV’, where the binding sites are indicated by underlined residues. The target tokenization for protein-ligand site prediction would then be represented as {2, 3, 5, 9}. Similarly, we follow the same strategy for tokenization of self-supervised pre-training tasks (Figure 2).

**Figure 2.**
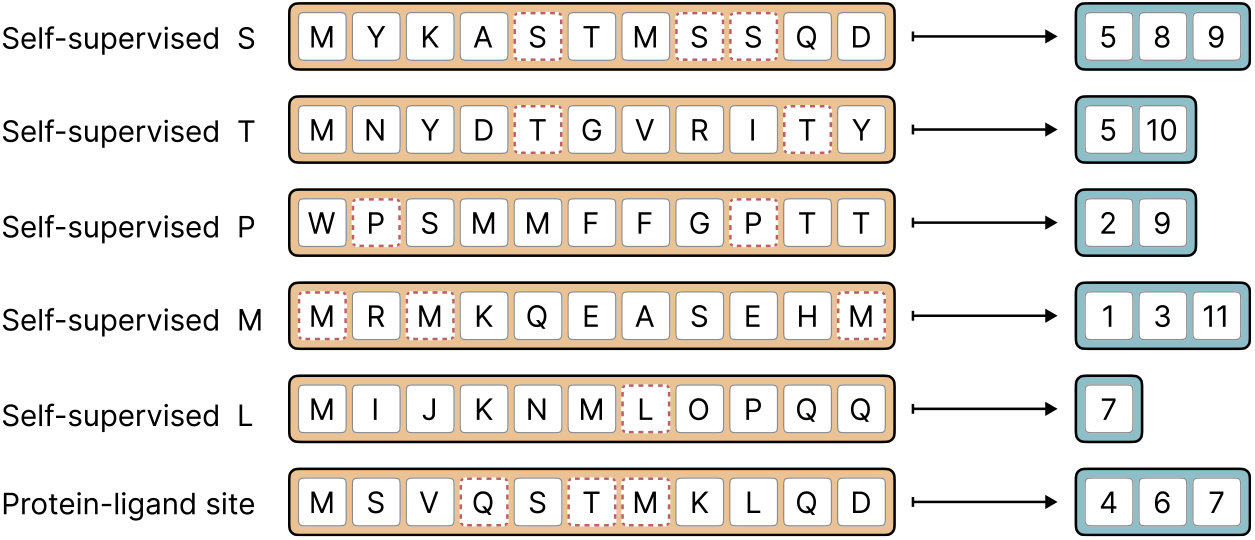
Tokenization of the self-supervised and protein-ligand binding site prediction tasks.

**Figure 3.**
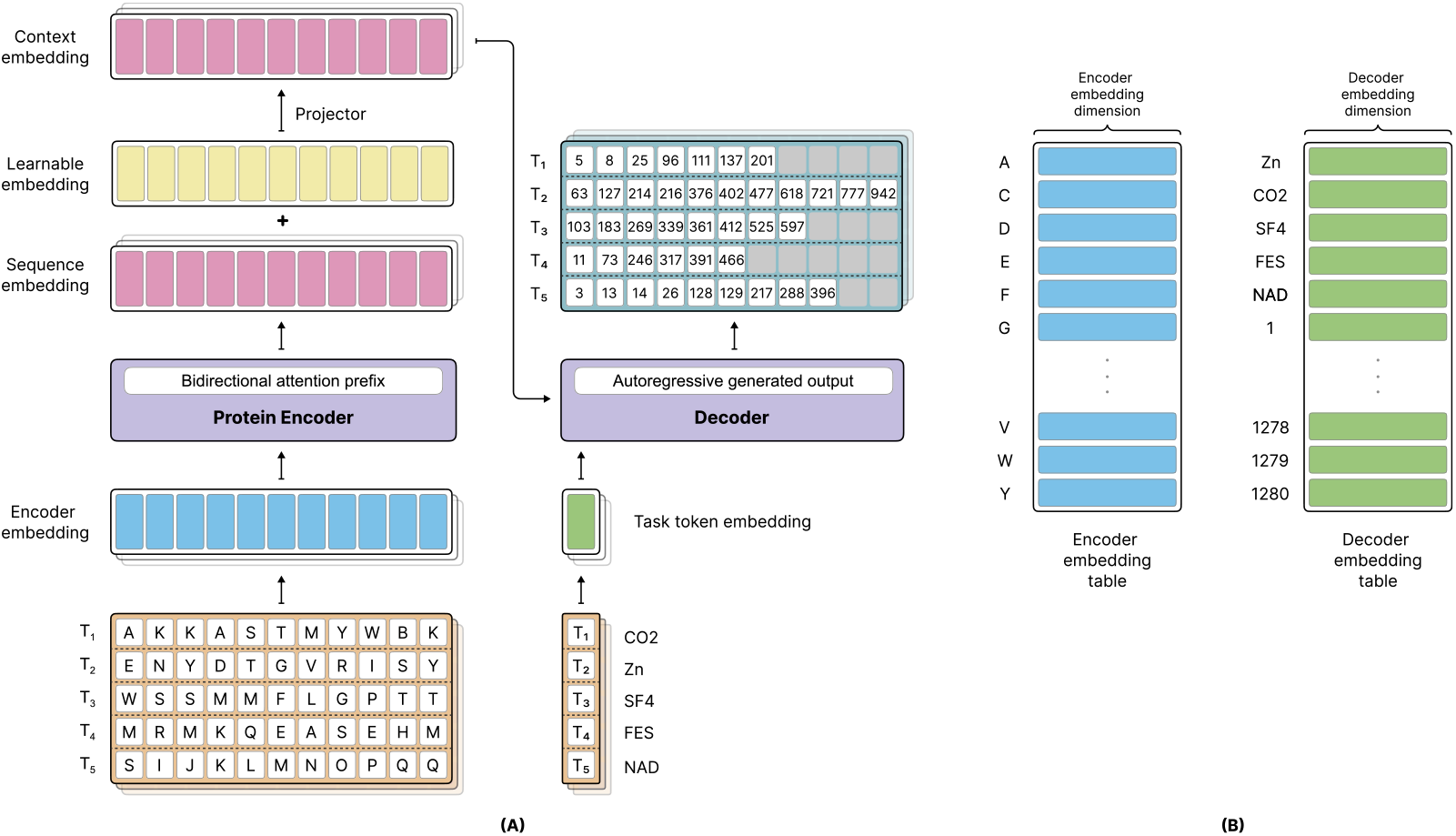
Inference flow of our framework for protein-ligand binding site prediction. (A) The model pipeline demonstrates the integration of a bidirectional protein encoder and a causal decoder. Protein sequences are processed into encoder embeddings, which then are combined with learnable positional and project down to be the context of the decoder via cross attention. Task-specific token embeddings guide the decoder to generate autoregressive outputs for binding site prediction. (B) The encoder and decoder utilize distinct embedding tables to represent amino acids in the encoder, and ligand-specific task tokens and position tokens in the decoder sides.

### Dataset

In this work, we consider the BioLip2 database Zhang et al. (2024a). We used the non-redundant dataset, which includes protein receptor sequences clustered at a 90% identity cutoff, along with annotations for each ligand-protein interaction site. To further enhance model robustness and increase the difficulty of the prediction task, we applied an additional clustering step at a 40% identity cutoff using CD-HIT Fu et al. (2012) to separate training, validation and test sets. To ensure a sufficient data ratio for training, evaluation, and testing, we included all ligands with more than 100 associated protein sequences, resulting in a total of 41 ligands. The details of the data preparation pipeline are provided in Appendix A.3.

In addition to the mentioned methods, we used a series of methods to interpret the task token embeddings which are elaborated in Appendix A.4

## 4 Experiments

In this section, we started with preparing the self-supervised pre-training of the decoder part. Then, we combined different protein-ligands tasks and trained a unified model with different number of ligands. After that, we scrutinized each ligand to find the correlation of learnable embeddings versus chemical properties to show global and local relationships between different ligands. For all our experiments, we considered ESM-2 650m model as the initialization of protein encoder of Prot2Token. In addition, we employed AdamW optimizer Loshchilov (2017) with weight decay of 0.1, and *β*_1_ = 0.9, *β*_2_ = 0.999, and epsilon to 1e-16 as the default hyperparameters of all experiments. Our learning rate strategy was based on cosine annealing with initial warm-up steps Loshchilov & Hutter (2016) starting from 1e-6 to 5e-5 for the first 256 steps, otherwise mentioned. All training experiments were developed using the PyTorch 2 framework Ansel et al. (2024) on one Nvidia 4×A100 80GB node.

### 4.1 Self-supervised pre-training

In the first step, we randomly sampled 4 million protein sequences from the UniRef50 database Suzek et al. (2015) for training and 4k for validation data. From them, we artificially created 80 million and 20k self-supervised samples, subsequently, by crafting each amino-acid-type/protein as one sample. Again, we sampled 1 million and 1k samples from them, respectively, to build the training and validation sets.

We used input sequence length of 1280, weight decay of 0.01 and batch size of 192 samples, equivalent to 73,728 tokens. Also, we set the warmup steps to 512. We only froze the encoder weights and made other parameters trainable. After training for 16 epochs, the model showed perplexity of 2.31 on the validation set. This indicates that it almost perfectly converted the embeddings out of encoder back to their original protein sequences.

### 4.2 Protein-ligand binding site

Based on the order of ligands in the table in Appendix A.4, we grouped the ligands into distinguishable sets of 10, 20, 30, and all 41 ligands. Each ligand in a set was treated as a separate task defined by a task token, training together. We selected each of those sets and jointly trained them alongside 20 self-supervised tasks using the latest checkpoint from the self-supervised pre-training phase. For this fine-tuning phase, the self-supervised tasks were reduced to a total number of 20k samples. Also, we removed protein samples with lengths greater than 1280 and set batch size to 98,304 tokens. During all training processes, only the last eight blocks of the encoder (ESM2-650m) were fine-tuned, while all non-encoder parameters of the supermodel were fully fine-tuned.

The effect of increasing the number of tasks for the first top 10 ligands is shown in Table 1. The detailed performance of all ligands is reported in Appendix A.4.

**Table 1:** F1 scores for the top 10 ligands across different training configurations on the test sets, with varying numbers of auxiliary ligands. The table summarizes the impact of jointly training with 10, 20, 30, and 41 ligands on binding site prediction. * indicates that pre-trained decoder weights were not used, and *†* indicates that self-supervised tasks were excluded during supervised training.

| Ligand | 10 ligands †* | 10 ligands* | 10 ligands | 20 ligands | 30 ligands | 41 ligands |
| --- | --- | --- | --- | --- | --- | --- |
| Zn<sup>2+</sup> | 0.0678 | 0.0657 | 0.7434 | 0.7498 | 0.7594 | 0.7575 |
| CA<sup>2+</sup> | 0.1022 | 0.0888 | 0.6566 | 0.6493 | 0.6472 | 0.6474 |
| CLA | 0.2749 | 0.2519 | 0.477 | 0.3763 | 0.4936 | 0.4762 |
| FAD | 0.1744 | 0.1476 | 0.6882 | 0.6565 | 0.6473 | 0.6537 |
| HEM | 0.243 | 0.232 | 0.6554 | 0.6698 | 0.6871 | 0.6796 |
| NAD | 0.1662 | 0.1248 | 0.6862 | 0.6851 | 0.6862 | 0.6952 |
| ADP | 0.1105 | 0.1053 | 0.6057 | 0.5779 | 0.5897 | 0.5834 |
| MG<sup>2+</sup> | 0.482 | 0.0326 | 0.4466 | 0.4603 | 0.4522 | 0.4575 |
| NAP | 0.1559 | 0.1417 | 0.6629 | 0.6813 | 0.6861 | 0.6746 |
| ATP | 0.1059 | 0.0927 | 0.4538 | 0.4355 | 0.5317 | 0.505 |
| **Average** | **0.1883** | **0.1283** | **0.6076** | **0.5942** | **0.6181** | **0.6132** |
| **Weighted Average** | **0.1849** | **0.0900** | **0.6297** | **0.6277** | **0.6368** | **0.6353** |

It is worth noting that while we could have excluded the self-supervised tasks entirely from fine-tuning stage, retaining a portion of these samples resulted in a measurable improvement in the model’s performance on the supervised protein-ligand tasks.

Direct comparison of our method with other available methods was not easy and straightforward due to several technical issues and potential overlap between their training data and our test sets; however, results of the comparison are provided in Appendix A.4.

### 4.3 Finding relationships between ligands

In this section, we scrutinized the task token embeddings of the decoder that pre-trained on all 41 ligands in the previous section to find the sign of chemical properties of ligands and their relationships together.

Empirically, based on the F1 scores of the ligands that the model was trained and evaluated on, the task token embeddings successfully captured meaningful representations of the ligands. However, to solidify this framework as a foundation for future research, we aimed to validate these embeddings from an additional perspective. Our goal is to create a robust infrastructure that can incorporate more ligands into a single model, thereby addressing the scarcity of data for certain ligands through knowledge transfer between ligands. To achieve this, first, from all 41 ligands, we selected top 28 ligands based on F1 score and filtered the rest and then, we analyzed the task token embeddings of remaining ligands by clustering them to explore ligand similarities in the embedding space.

Simultaneously, we clustered the ligands based on their biochemical features in the real world and in the last step, we investigated the correlation between these two clustering approaches. The purpose of this comparison was to determine whether the learned task token embeddings genuinely reflect real-world relationships between ligands or if they merely memorize specific patterns without capturing meaningful biochemical similarities. Figure 4 highlights the intersection between the two spaces of ligand representations: the embedding space and the biochemical feature space. It illustrates which ligands or sets of ligands have their relationships successfully captured by the generated task token embeddings, as reflected by their agreement with relationships derived from biochemical features, and which embeddings failed to capture such relationships. More details including feature selection, methods and the interpretation algorithm are placed in Appendix A.4.

**Figure 4.**
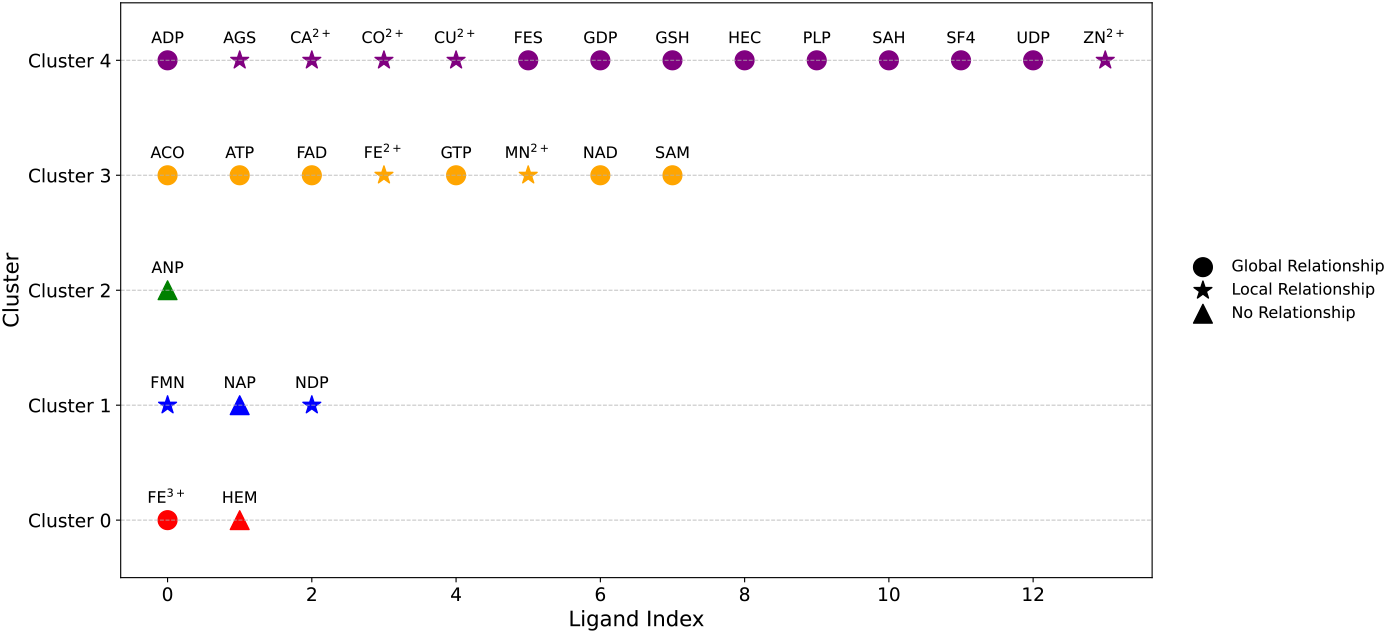
Global Relationships indicate that general biochemical features shared among many ligands have been captured. Local Relationships reflect the successful capture of biochemical properties between specific ligands and their closely related counterparts. No Relationships indicate that the biochemical properties were not captured at all.

### 4.4 Multi-task learning effect

In this section, we tried to investigate the effect of multi-task learning on three ligands that have low number of samples but share similar semanticity in the embedding space of task tokens (Figure 4). Therefore, we selected GSH, CO, AGS as our target ligands because (1) they belong to the same cluster, (2) showed global or local relationships and, (3) have less than 200 or lower number of protein sequences in total. We considered three groups to measure the performance of three ligands. There groups are defined as follow:

**Group 1**. Combination of the target ligands, GSH, CO and AGS (Equivalent to 1,819 tokens).

**Group 2**. Combination of CLA, FAD, HEM and NAD as ligands that did not share a close semantic representation in Figure 4 (Equivalent to ∼ 43*k* tokens).

**Group 3**. Combination of Zn, Ca, ADP, members from cluster 4 (Equivalent to ∼ 37*k* tokens).

Need to point out, in order to make the comparison of group 2 and 3 fair, we considered the total number of tokens in these groups close to each other. Table 2 shows that group 3 which shares a similar cluster with the target ligands, improves F1 score more than other groups.

**Table 2:**
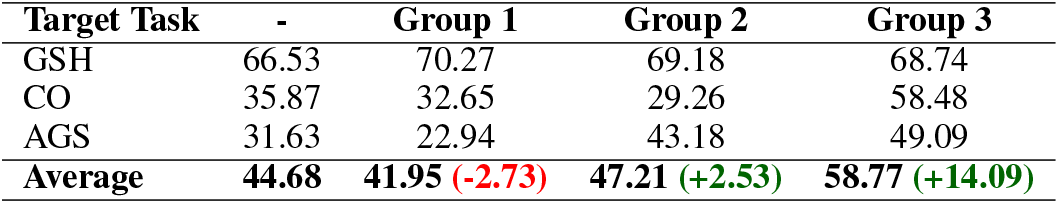
The effect of jointly training under representative ligands based on different auxiliary groups with respect to F1 score. “-” means no auxiliary task is used during training the target task.

### 4.5 Discussion

In this section, we describe our results in multiple parts, highlighting key findings and their implications. Each one addresses specific aspects of our unified framework for protein-ligand binding site prediction, including our approach’s generality, interpretability, and future improvements.

#### 4.5.1 A unified protein-ligand binding sites prediction

Our method demonstrates strong predictive performance across various ligands, achieving competitive F1 scores, particularly for those with sufficient training samples. Self-supervised pre-training on a large corpus of protein sequences addresses an important challenge of the Prot2Token architecture by equipping the decoder with implicit inductive biases, injecting effective knowledge for learning for binding site prediction. Additionally, multi-task learning with a lot of tokens plays a crucial role in addressing data scarcity and improving the average performance. Training on multiple ligands simultaneously allows the model to leverage shared information, creating a synergistic effect where learning one ligand enhances predictions for others (Table 1).

#### 4.5.2 Interpretation

The interpretation of task token embeddings highlights the model’s ability to capture meaningful relationships among ligands, revealing global and local similarities that align with their biochemical properties. By classifying relationships into global, local, and no relationships (Figure 4), we identified both the strengths and limitations of these embeddings. While the embeddings effectively captured true relationships for many ligands, they struggled for others, indicating areas for further refinement. To validate the utility of these embeddings, we correlated biochemical feature-based clusters with embedding-based clusters, demonstrating a strong alignment for most ligands. This validation underscores the capacity of the embeddings to encode biologically relevant information and supports their extension to new ligands. Moreover, leveraging these relationships through auxiliary tasks in multi-task learning significantly improved predictions for underrepresented ligands (Table 2). Our results correlate this to the existence of a high number of auxiliary tokens plus the similarity of embeddings. However, the presence of no relationships suggests either limitations in the embeddings themselves or gaps in the biochemical features used, pointing to opportunities for better interpretation. These findings illustrate how interpretation-driven insights can guide multi-task learning strategies, ultimately enhancing predictions for heavily underrepresented ligands.

#### 4.5.3 Future work

While the proposed framework demonstrates strong predictive performance and generality across diverse ligands, several avenues remain for future exploration to enhance its robustness and applicability. One promising direction is the use of pre-trained chemical encoders to represent ligands, replacing the current learnable task token approach. This could allow the model to leverage rich, pre-existing knowledge about chemical properties, potentially improving predictions for ligands with limited training data. Another critical extension involves incorporating 3D structural information of proteins as contextual input. The inclusion of spatial features could enable the model to better capture the intricate relationships between protein residues and ligands.

Additionally, performance variability across ligands highlights persistent challenges, including prediction complexity and data scarcity, particularly for underrepresented ligands. While our approach significantly improves performance through techniques like multi-task learning, the model’s accuracy remains dependent on the availability of larger and more diverse datasets. Expanding our framework to include more comprehensive datasets and advanced ligand representations holds significant potential for further advancements in protein-ligand binding site prediction.

### Meaningfulness Statement

Understanding life at a molecular level requires identifying how proteins interact with various ligands, a fundamental process in biological functions such as metabolism, signaling, and disease progression. A meaningful representation of life should capture these interactions with accuracy, enabling deeper insights into cellular mechanisms and facilitating drug discovery. Our work contributes to this direction by leveraging autoregressive transformer models to predict protein-ligand binding sites directly from protein sequences. This approach enhances interpretability and generality in protein-ligand interaction prediction, paving the way for more effective computational tools that aid in biomedical research and therapeutic development.

A Appendix

### A.1 Related work

We categorized the development of diverse methodologies for protein binding site prediction into two main clusters. Standalone models, which propose entirely new methods, and ensemble models, which integrate existing techniques to enhance performance.

#### A.1.1 Standalone models

These approaches introduce novel methods to address the binding site prediction problem and can be further divided into three subcategories based on their approaches to feature extraction and processing:

1. Classical Approaches, which rely on traditional feature extraction techniques such as sequence-based and structure-based features.
2. Deep Learning Approaches, which utilize advanced architectures like graph neural networks (GNNs), ResNet, and BiLSTM to automatically extract high-level features from protein data.
3. Large Language Model-Based Approaches, which employ state-of-the-art language models trained on massive protein sequence datasets to capture complex patterns and long-range dependencies.

##### Classical Approaches

Yu et al. (2013) developed TargetS, a tool designed to predict binding sites for twelve ligand types. TargetS combines protein evolutionary information, predicted secondary structure, and ligand-specific binding propensities of residues to construct discriminative features. To address the imbalance between binding and non-binding residues, it employs random undersampling and classifier ensembles. Additionally, TargetS uses MODELLER, a software package for predicting 3D structures from protein sequences, to perform spatial clustering of the predicted binding residues and identify the final binding sites. Yang et al. (2013) introduced the COACH method, which combines TM-SITE and S-SITE to provide accurate predictions of binding sites. TM-SITE utilizes the query protein structure to identify relevant templates through structural alignment. These alignments are then used to identify potential binding sites, which are further refined by clustering the ligand centers of the templates to predict the final binding sites. S-SITE, on the other hand, uses the query sequence to generate a frequency profile that captures evolutionary information by aligning the query sequence with similar sequences from a database. Relevant templates are then identified, and aggregation with a voting scheme is applied to determine the binding residues.

##### Deep Learning Approaches

Xia et al. (2020) presented DELIA, a deep-learning-based method for predicting protein–ligand interactions across five ligand types. The method concatenates extracted sequence-based features and combines them with a 2D structure-based distance map. This distance map is generated by calculating the distances between each pair of residues in the protein structure, after which ResNet is used to extract spatial features from the map. A fully connected layer is then applied to merge both feature types into a single vector representation. To address the issue of data imbalance, DELIA employs random undersampling by selecting 20% of the negative samples along with all positive samples in each subset. Additionally, oversampling is applied in mini-batches to ensure adequate representation of samples during training. Essien et al. (2019) introduce ZinCaps, which utilize Capsule Networks to predict zinc ion binding sites using only protein sequence data. The Capsule Networks dynamically adjust and refine the contributions of lower-level features, resulting in better high-level feature representations. ZinCaps employs a sliding window of 25 residues around each target residue to extract features. Xia et al. (2021) utilize Hierarchical Graph Neural Networks (HGNNs) to represent protein structures by modeling residues as nodes within a graph. The HGNN framework consists of three key modules: a GNN-Encoder, which encodes raw feature vectors into high-level representations; GNN-Blocks, which extract hierarchical features by updating node, edge, and graph-level information; and a multi-layer perceptron (MLP) classifier, which predicts nucleic-acid-binding residues, specifically targeting DNA- and RNA-binding residues. Cui et al. (2019) utilize a deep convolutional neural network for binding site prediction involving fourteen ligands of metal ions and small molecules. Seven residue-level features are extracted, forming a matrix based on sequence length and features, which is then passed to the encoder. The encoder extracts high-level feature representations from this input matrix. During training, the decoder leverages the initial predictions, combines them with the encoded features, and feeds them back into the decoder to generate the final predictions. Essien et al. (2025) proposed GPred, a method that utilizes geometry-aware GNNs to predict binding sites for seven small metal ion ligands. Protein structures are converted into 3D point clouds, where each point represents an atom, which is associated with a 59-dimensional feature vector comprising physicochemical properties and evolutionary information. A ball query strategy is applied to define a local graph for each atom, which is then processed by a Point Transformer to learn structural and biochemical features of the protein. Finally, a fully connected layer is used for classification.

##### Large Language Models

Yuan et al. (2022) introduced LMetalSite, a framework that uses ProtTrans, a pretrained transformer model, to extract protein sequence embeddings. The model leverages multi-task learning to improve predictive performance by utilizing shared information across different tasks. It employs a stack of transformer encoder layers as a shared network to capture common characteristics, such as long-range dependencies. Finally, four ion-specific MLPs are used to learn the binding patterns of individual metal ions. Essien et al. (2023) proposed IonPred, a model based on ELECTRA, a transformer architecture with two components: a generator, which is considered successful when it generates replacements that are difficult for the discriminator to identify, and a discriminator, which is considered successful when it accurately identifies the replaced tokens generated by the generator. ELECTRA creates embeddings by converting tokens into dense vectors, which are subsequently refined using transformer-based generator and discriminator networks to produce contextualized embeddings. These embeddings serve as the foundation for IonPred’s predictions.

#### A.1.2 Ensemble methods

These approaches integrate pre-existing models and ensemble their predictions to achieve improved performance. Xia et al. (2020) developed BindWeb, a web server that uses the GraphBind and DELIA models to predict potential binding residues and clusters them spatially to identify distinct binding pockets on the protein surface. BindWeb supports predictions for DNA, RNA, and five other ligands, integrating results from both models and applying mean shift clustering to determine binding pockets. Hu et al. (2016) introduced two methods, IonSeq and IonCom, for predicting binding sites of 13 metal ions and acid radical ion ligands. IonSeq is a sequence-based method that uses sequence profiles and a modified AdaBoost algorithm to balance predictions between binding and non-binding residues. IonCom Hu et al. (2016) is a ensemble method that integrates IonSeq with multiple template-based approaches (e.g., COFACTOR, TM-SITE, S-SITE, and COACH) to enhance prediction accuracy and robustness. ATPbind Hu et al. (2018) integrated the outputs of two template-based predictors (S-SITE and TM-SITE) and employs an undersampling technique to address data imbalance. Multiple SVM models are trained, and their outputs are combined using the mean ensemble method to produce the final predictions. Lin et al. (2016) developed the MIB web server, which predicts binding sites for 12 types of metal ions by aligning the query protein with templates and assigning binding scores based on sequence and structural similarity without relying on machine learning models or training.

### A.2 Architecture

The proposed architecture integrates an encoder-decoder framework, where each component plays a distinct role in processing and generating sequences. The encoder, denoted as *f*_enc_(·), processes the input sequence of tokens *x* = (*x*_1_, *x*_2_, …, *x*_*N*_) to produce contextual embeddings *h*_enc_:

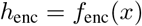

Where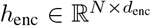, *N* is the sequence length, and *d*_*enc*_ is the encoder’s hidden dimension. To enhance the contextual embeddings from the encoder, a learnable embedding layer *g*_pos_(·) is introduced, which adds positional information to the encoder’s output. The final encoder representation *h*_*aug*_ is obtained by summing the positional embeddings p with the encoder output:

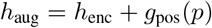

where 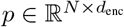 is the learnable positional embedding. Since the decoder operates at a potentially different hidden dimension *d*_*dec*_ a linear projection layer 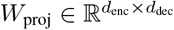 is applied to align the encoder’s output to the decoder’s dimensionality:

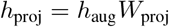

where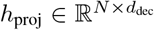. The decoder, *f*_dec_(·), receives two inputs: the projected encoder output *h*_*proj*_and a sequence of task tokens *t* = (*T*_1_, *T*_2_, …, *T*_*m*_). Task tokens are mapped to their respective embeddings via a learnable embedding layer:

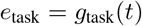

where 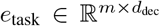. Finally, the decoder attends to both the encoder’s projected output using crossattention and, the task token embeddings, producing the sequence of predictions *y* = (*y*_1_, *y*_2_, …, *y*_*N*− *M*_). The decoder is formulated as:

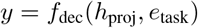

where the decoder predicts all non-task tokens conditioned on the projected encoder output and task token embeddings.

The encoder portion of the model follows the exact architecture of the ESM-2 650.m model. Its output is first combined with a learnable embedding, then projected from 1280 to 640 dimensions using a learnable linear projector. The decoder is a standard causal (autoregressive) Transformer with a hidden size of 640, a feed-forward dimension of 2560, GeLU activations and 16 attention heads, which iterated over 16 blocks. Additionally, FlashAttention.2 Dao (2023) is employed to improve both training speed and memory efficiency.

### A.3 Datasets

BioLip2 is one of the most comprehensive databases for ligand-protein interactions, primarily derived from the Protein Data Bank (PDB). Each entry in BioLip2 includes detailed annotations on ligand-binding residues, ligand-binding affinities, catalytic site residues, enzyme commission (EC) numbers and gene ontology (GO) terms. The database is also cross-linked with external resources, including RCSB PDB, UniProt, and PubMed. To obtain protein sequences, we used receptor sequences clustered at a 90% identity cutoff. For annotations, we retrieved data for each ligand-protein interaction site. To increase the complexity of binding site prediction and enhance model robustness, we further clustered the data at a 40% identity cutoff. This additional clustering step helps prevent data leakage between training, evaluation, and testing datasets. We first removed DNA and RNA sequences and excluded any sequences with fewer than 50 residues. Next, we generated FASTA files containing residues and annotations for all 5,717 ligands. We then applied a threshold cutoff, selecting ligands that bind with over 100 sequences, resulting in 41 ligands. We aimed to balance selecting the most significant ligands based on a literature review while ensuring a sufficient number of samples for training and testing the model. We used CD-HIT to cluster the data with a 40% identity cutoff before splitting the data into training, evaluation, and testing datasets. Because of the limited number of samples and to ensure sufficient data for testing, we used two splitting ratios: 70%, 10%, and 20% for training, evaluation, and testing, respectively, for the first 30 ligands in Table 3, and also, 50%, 20%, and 30% for training, evaluation, and testing, respectively, for the remaining ligands.

**Table 3:** Dataset statistics of all ligands.

| Ligand No | Chemical Formula | Name | BioLip Fasta Name | Num Sequences | Binding Sites |
| --- | --- | --- | --- | --- | --- |
| 1 | Zn<sup>2+</sup> | Zinc Ion | ZN.fasta | 4665 | 23310 |
| 2 | Ca<sup>2+</sup> | Calcium Ion | CA.fasta | 3043 | 22161 |
| 3 | CLA | Chlorophyll A | CLA.fasta | 342 | 17690 |
| 4 | FAD | Flavin-Adenine Dinucleotide | FAD.fasta | 825 | 16583 |
| 5 | HEM | Heme | HEM.fasta | 845 | 13118 |
| 6 | NAD | Nicotinamide Adenine Dinucleotide | NAD.fasta | 658 | 10615 |
| 7 | ADP | Adenosine Diphosphate | ADP.fasta | 941 | 9899 |
| 8 | MG<sup>2+</sup> | Magnesium Ion | MG.fasta | 2951 | 9494 |
| 9 | NAP | Nicotinamide Adenine Dinucleotide Phosphate | NAP.fasta | 462 | 8108 |
| 10 | ATP | Adenosine Triphosphate | ATP.fasta | 680 | 7635 |
| 11 | HEC | Heme C | HEC.fasta | 264 | 7296 |
| 12 | SF4 | Iron/Sulfur Cluster | SF4.fasta | 509 | 5834 |
| 13 | FMN | Flavin Mononucleotide | FMN.fasta | 437 | 5789 |
| 14 | SAH | S-Adenosyl-L-Homocysteine | SAH.fasta | 392 | 4675 |
| 15 | NDP | Nucleotide Diphosphate | NDP.fasta | 243 | 4301 |
| 16 | ANP | Adenylyl-imidodiphosphate | ANP.fasta | 354 | 3861 |
| 17 | GDP | Guanosine Diphosphate | GDP.fasta | 339 | 3792 |
| 18 | GLC | Glucose | GLC.fasta | 454 | 3674 |
| 19 | PLP | Pyridoxal-5'-Phosphate | PLP.fasta | 377 | 3608 |
| 20 | MN<sup>2+</sup> | Manganese Ion | MN.fasta | 789 | 3315 |
| 21 | COA | Coenzyme A | COA.fasta | 259 | 2870 |
| 22 | SAM | S-Adenosylmethionine | SAM.fasta | 214 | 2540 |
| 23 | AMP | Adenosine Monophosphate | AMP.fasta | 275 | 2430 |
| 24 | BGC | Beta-D-Glucose | BGC.fasta | 331 | 2375 |
| 25 | FE<sup>3+</sup> | Ferric Ion | FE.fasta | 532 | 2268 |
| 26 | MAN | Mannose | MAN.fasta | 446 | 2047 |
| 27 | FES | Iron-Sulfur Cluster | FES.fasta | 272 | 1986 |
| 28 | PO<sub>4</sub><sup>3-</sup> | Phosphate Ion | PO4.fasta | 378 | 1908 |
| 29 | GTP | Guanosine Triphosphate | GTP.fasta | 150 | 1724 |
| 30 | UDP | Uridine Diphosphate | UDP.fasta | 154 | 1601 |
| 31 | CU<sup>2+</sup> | Copper Ion | CU.fasta | 331 | 1530 |
| 32 | GSH | Glutathione | GSH.fasta | 200 | 1516 |
| 33 | AGS | Agmatine Sulfate | AGS.fasta | 136 | 1512 |
| 34 | ACO | Aconitase | ACO.fasta | 108 | 1482 |
| 35 | GAL | Galactose | GAL.fasta | 233 | 1188 |
| 36 | SO<sub>4</sub><sup>2-</sup> | Sulfate Ion | SO4.fasta | 218 | 1177 |
| 37 | CLR | Cholesterol | CLR.fasta | 176 | 1112 |
| 38 | Y01 | Cholesterol Hemisuccinate | Y01.fasta | 106 | 991 |
| 39 | BMA | Beta-Mannose | BMA.fasta | 158 | 696 |
| 40 | FE<sup>2+</sup> | Ferrous Ion | FE2.fasta | 186 | 675 |
| 41 | CO<sup>2+</sup> | Cobalt Ion | CO.fasta | 160 | 660 |

### A.4 Experiments

For fine-tuning on the protein-ligand datasets, the model was trained on a combined training set of selected ligands. During training, validation was performed for each ligand individually, and the best checkpoint for each ligand was saved based on its validation set performance. At the end of training, these best checkpoints were evaluated on their respective test sets. Figure 5 shows the average validation F1 score across epochs, with the highest average performance observed at epoch 30. However, this checkpoint showed slightly lower average test performance compared to using individual best checkpoints for each ligand.

**Figure 5.**
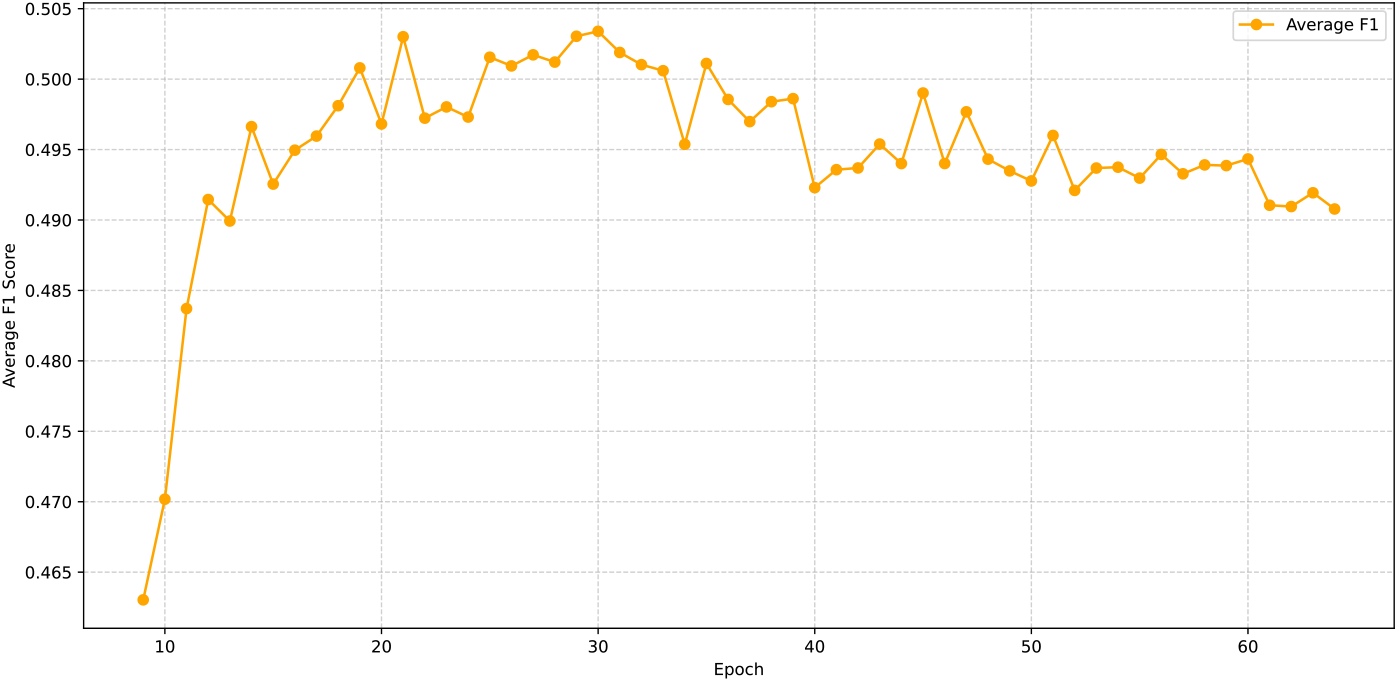
Average of F1 values for all 41 ligands during the training based on the validation sets. The performance peaked at epoch 30.

The results for all ligands are presented in Table 4. To compute the metric for the autoregressive model’s output, each amino acid in a protein was treated as an individual positive or negative sample. Predicted binding residues from the decoder were considered positive samples, while all other amino acids were treated as negative (zero) samples. The metrics were then calculated based on this classification.

**Table 4:** F1 scores of positive labels for all ligands across different training configurations, with varying numbers of auxiliary ligands on the test sets. The table summarizes the impact of jointly training with 10, 20, 30, and 41 ligands on binding site prediction. * indicates that pre-trained decoder weights were not used, and † indicates that self-supervised tasks were excluded during supervised training.

| Ligands | 10 tasks †* | 10 tasks* | 10 tasks | 20 tasks | 30 tasks | 41 tasks |
| --- | --- | --- | --- | --- | --- | --- |
| ZN<sup>2+</sup> | 0.0678 | 0.0657 | 0.7434 | 0.7498 | 0.7594 | 0.7575 |
| CO<sup>2+</sup> | 0.1022 | 0.0888 | 0.6566 | 0.6493 | 0.6472 | 0.6474 |
| CLA | 0.2749 | 0.2519 | 0.477 | 0.3763 | 0.4936 | 0.4762 |
| FAD | 0.1744 | 0.1476 | 0.6882 | 0.6565 | 0.6473 | 0.6537 |
| HEM | 0.243 | 0.232 | 0.6554 | 0.6698 | 0.6871 | 0.6796 |
| NAD | 0.1662 | 0.1248 | 0.6862 | 0.6851 | 0.6862 | 0.6952 |
| ADP | 0.1105 | 0.1053 | 0.6057 | 0.5779 | 0.5897 | 0.5834 |
| MG<sup>2+</sup> | 0.482 | 0.0326 | 0.4466 | 0.4603 | 0.4522 | 0.4575 |
| NAP | 0.1559 | 0.1417 | 0.6629 | 0.6813 | 0.6861 | 0.6746 |
| ATP | 0.1059 | 0.0927 | 0.4538 | 0.4355 | 0.5317 | 0.505 |
| **Average (top 10)** | **0.1883** | **0.1283** | **0.6076** | **0.5942** | **0.6181** | **0.6130** |
| HEC | - | - | - | 0.6438 | 0.6511 | 0.6537 |
| SF4 | - | - | - | 0.6508 | 0.584 | 0.5685 |
| FMN | - | - | - | 0.6921 | 0.6983 | 0.6945 |
| SAH | - | - | - | 0.6385 | 0.6473 | 0.6503 |
| NDP | - | - | - | 0.7122 | 0.7085 | 0.6979 |
| ANP | - | - | - | 0.6153 | 0.6214 | 0.6217 |
| GDP | - | - | - | 0.5948 | 0.6335 | 0.6465 |
| GLC | - | - | - | 0.2091 | 0.2237 | 0.2214 |
| PLP | - | - | - | 0.777 | 0.778 | 0.762 |
| MN<sup>2+</sup> | - | - | - | 0.7278 | 0.7245 | 0.7376 |
| **Average (top 20)** | - | - | - | **0.6102** | **0.6172** | **0.6130** |
| COA | - | - | - | - | 0.3978 | 0.4011 |
| SAM | - | - | - | - | 0.6355 | 0.6252 |
| AMP | - | - | - | - | 0.4316 | 0.4432 |
| BGC | - | - | - | - | 0.2165 | 0.1932 |
| FE<sup>3+</sup> | - | - | - | - | 0.6756 | 0.6606 |
| MAN | - | - | - | - | 0.1407 | 0.1216 |
| FES | - | - | - | - | 0.7162 | 0.7018 |
| PO<sub>4</sub><sup>3-</sup> | - | - | - | - | 0.2288 | 0.2278 |
| GTP | - | - | - | - | 0.5332 | 0.5461 |
| UDP | - | - | - | - | 0.5522 | 0.5391 |
| **Average (top 30)** | - | - | - | - | **0.566** | **0.5615** |
| CU<sup>2+</sup> | - | - | - | - | - | 0.5607 |
| GSH | - | - | - | - | - | 0.6924 |
| AGS | - | - | - | - | - | 0.5301 |
| ACO | - | - | - | - | - | 0.5026 |
| GAL | - | - | - | - | - | 0.2762 |
| SO<sub>4</sub><sup>2-</sup> | - | - | - | - | - | 0.1386 |
| CLR | - | - | - | - | - | 0.0373 |
| Y01 | - | - | - | - | - | 0.0419 |
| BMA | - | - | - | - | - | 0.2273 |
| FE<sup>2+</sup> | - | - | - | - | - | 0.6033 |
| CO<sup>2+</sup> | - | - | - | - | - | 0.517 |
| **Average (all)** | - | - | - | - | - | **0.5115** |

To provide a comparison of our model’s performance with other available methods, we present the results in Table 5. However, the comparison process faced several challenges: some web servers were not operational during testing, while others only allowed predictions on individual samples, making bulk evaluation difficult and very slow to response. We attempted to evaluate IonCom, and MIB2 [33] server tools, but encountered several issues: MIB2 had extremely slow response times, and IonCom imposed strict sample limitations for evaluation. Additionally, a potential overlap between the training data of these methods and our crafted test sets further made a fair evaluation complicated. This was particularly evident for LMetalSite, where their reported performance on their own test sets was significantly lower compared to their results on our test sets, indicating the sign of data leaking in this comparison.

**Table 5:** Comparison of our method’s best performance for each ligand with other available methods on selected ligands based on F1 score. The main values are based on their reported test set performance as described in their respective papers. * Indicates they are reported on our test sets.

| Ligand | Metrics | Prot2Token<br>(Our method) | TargetS<br>Yu et al. (2013) | LMetalSite<br>Yuan et al. (2022) | ZinCap<br>Essien et al. (2019) | MIB2<br>Lu et al. (2022) |
| --- | --- | --- | --- | --- | --- | --- |
| CA<sup>2+</sup> | F1 | 0.6566* | 0.392* | 0.526 (0.7370*) | - | - |
|  | MCC | - | 0.320 (0.431*) | 0.542 (0.7342*) | - | - |
|  | Acc | - | 0.984 (0.977*) | 0.9884* | - | 0.941 |
| MG<sup>2+</sup> | F1 | 0.4603* | 0.433* | 0.367 (0.5560*) | - | - |
|  | MCC | - | 0.383 (0.450*) | 0.419 (0.5773*) | - | - |
|  | Acc | - | 0.990 (0.992*) | 0.9949* | - | 0.946 |
| ZN<sup>2+</sup> | F1 | 0.7594* | 0.660* | 0.76 (0.8299*) | 0.451* | - |
|  | MCC | - | 0.557 (0.660*) | 0.761 (0.8275*) | 0.54 (0.48*) | - |
|  | Acc | - | 0.989 (0.989*) | 0.9953* | 0.870 (0.97*) | 0.948 |
| MN<sup>2+</sup> | F1 | 0.7376* | 0.579* | 0.662 (0.8048*) | - | - |
|  | MCC | - | 0.445 (0.574*) | 0.661 (0.8024*) | - | - |
|  | Acc | - | 0.987 (0.989*) | 0.995* | - | 0.950 |

#### A.4.1 Interpretation

In this study, we developed a protein binding site prediction model using a multi-task learning framework, where each task represents a specific ligand. A 640-dimensional task token was incorporated for each ligand alongside the protein sequences. During training, the model learned meaningful task token embeddings that effectively represent ligands and their unique characteristics.

To validate the task token embeddings, we employed two clustering approaches: one based on the trained task token embeddings and the other on biochemical ligand features. For precise clustering and clearer analysis, ligands with an F1 score below 0.5 were excluded to minimize noise, leaving 28 out of 41 ligands for analysis. Task token embeddings were reduced to 27 principal components using PCA, preserving 99% of the variance, and clustered with k-means to generate target clusters. For validating all ligands, the full set of 41 ligands was included. In this case, task token embeddings were reduced to 40 components to preserve 99% of the variance, and the same clustering method was applied. For the ligand features, 26 biochemical descriptors were collected, covering physical, chemical, electronic, hydrophilic, lipophilic, and geometric properties.

A systematic feature selection process evaluated all possible combinations of up to 13 features selected from these 26 descriptors (approximately 39 million combinations) to optimize clustering quality against the target clusters. The Adjusted Rand Index (ARI) was used as the selection metric, while Normalized Mutual Information (NMI) and Pairwise Accuracy metrics were later employed to evaluate the final selection.

The clustering results demonstrate that the learned task token embeddings are meaningful, as their clustering aligns closely with that based on ligand-specific biochemical features. Moderate-to-high agreement metrics (ARI = 0.447, NMI = 0.614, and pairwise accuracy = 0.783) highlight the embeddings’ ability to capture key biochemical characteristics of ligands. Chemically significant features, such as ‘MolecularWeight,’ ‘NetCharge,’ and ‘RotatableBonds,’ identified as part of the optimal feature set, further reinforce the relevance of the embeddings. The overlap and similarity in ligand grouping across both clustering approaches validate the hypothesis that the task token embeddings effectively encode biologically and chemically meaningful information.

However, reducing task token embeddings or biochemical features to 2D for visualization causes significant information loss, making 2D clustering plots less informative (Figures 11 and 12). These findings emphasize the importance of preserving higher-dimensional information for accurate interpretation and highlight the value of task token embeddings in ligand characterization for protein binding site prediction. Figure 6 shows the embeddings-based clustering, while Figure 7 shows the features-based clustering, and Figure 4 illustrates the global, local, and no relationships between the two approaches of embeddings-based clustering and features-based clustering.

**Figure 6.**
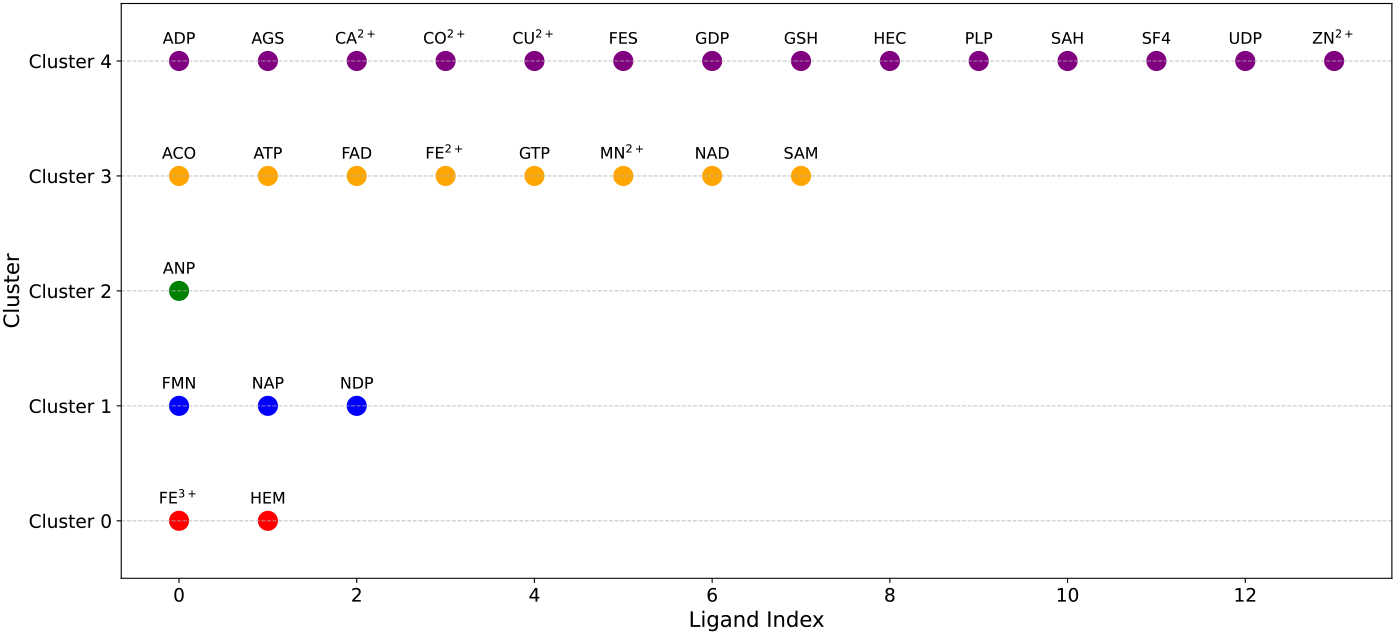
Clustering results of embeddings on top 28 ligands based on F1 score.

**Figure 7.**
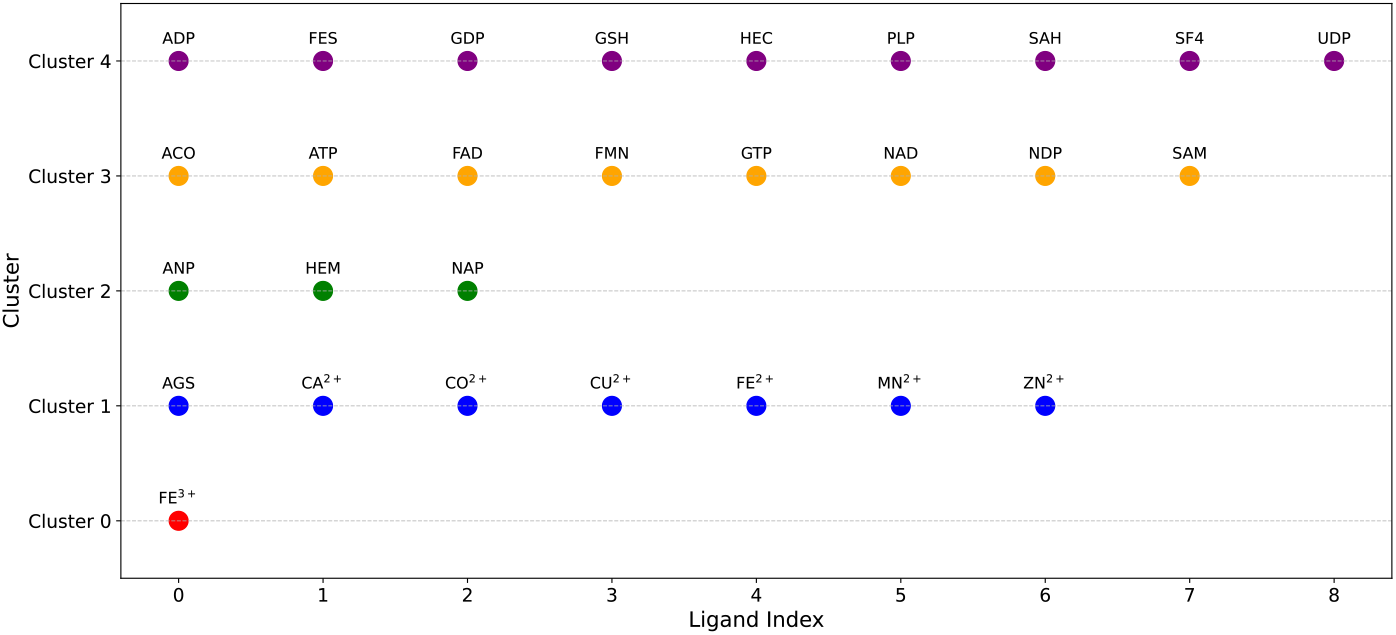
Clustering results of features on the 28 selected ligands.

##### Global relationships

Figure 4 highlights the ligands that have been clustered correctly across and within both clustering approaches. For instance, in Cluster 3, the solid circles for ACO, ATP, FAD, GTP, NAD, and SAM ligands represent ligands that have been consistently clustered across and within the same clusters in both approaches. This indicates that the task token embeddings successfully capture their similarity with each other and with the rest of the ligands.

##### Local relationships

Figure 4 also depicts ligands that have been clustered correctly only within clusters in both clustering approaches. For example, the stars in cluster 3 for FE^2+^ and MN^2+^ indicate that these ligands are grouped but appear in different clusters across the two approaches. Nevertheless, the task token embeddings still manage to capture their similarity with each other, even if they fail to capture their similarity with other ligands.

##### No relationships

For some ligands, the task token embeddings fail to accurately capture their global or local relationships. This may be due to the ligand features collected not being entirely representative and requiring further refinement, or because the task token embeddings themselves need improvement. Figure 4 illustrates these “no relationships” using triangles; for instance, the HEM ligand has been grouped with different ligands across different clusters in both approaches.

For further investigation of the task token embeddings, we incorporated all 41 ligands into the clustering analysis. The metrics showed a notable drop: *ARI* = 0.259, *NMI* = 0.333, and *PairwiseAccuracy* = 0.733. This decrease was expected, as including task token embeddings for ligands with low F1 scores introduced some misaligned clusters. However, a closer examination reveals that the embeddings still effectively capture the global and local relationships between most ligands. Figures 8 and 9 depict the embeddings-based clustering and features-based clustering, respectively, while Figure 10 illustrates the global, local, and no relationships across all 41 ligands. Notably, out of the 41 ligands, the task token embeddings successfully represented 21 ligands globally, 13 ligands locally, and misrepresented 7 ligands. These results indicate that the task token embeddings consistently demonstrate strong global and local relationships, effectively capturing biochemical similarities among ligands. This reinforces the conclusion that the model has learned meaningful representations, even for ligands with low F1 scores.

**Figure 8.**
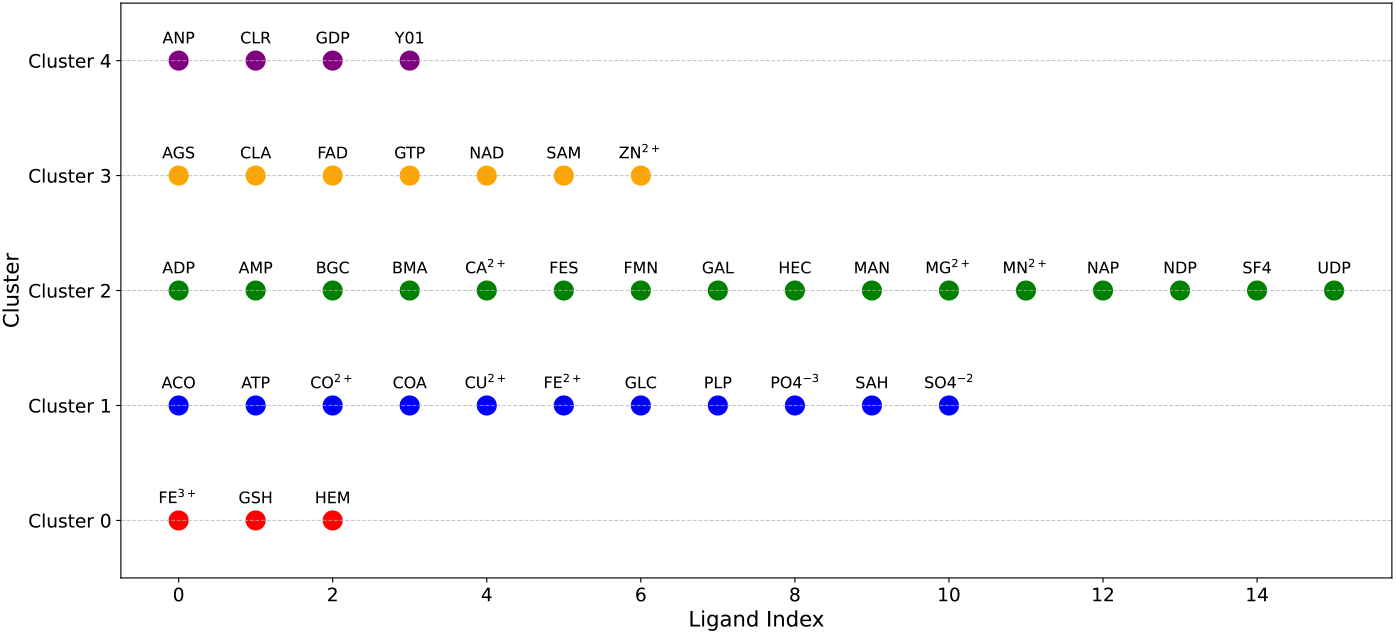
Clustering results of embeddings on all 41 ligands based on F1 score.

**Figure 9.**
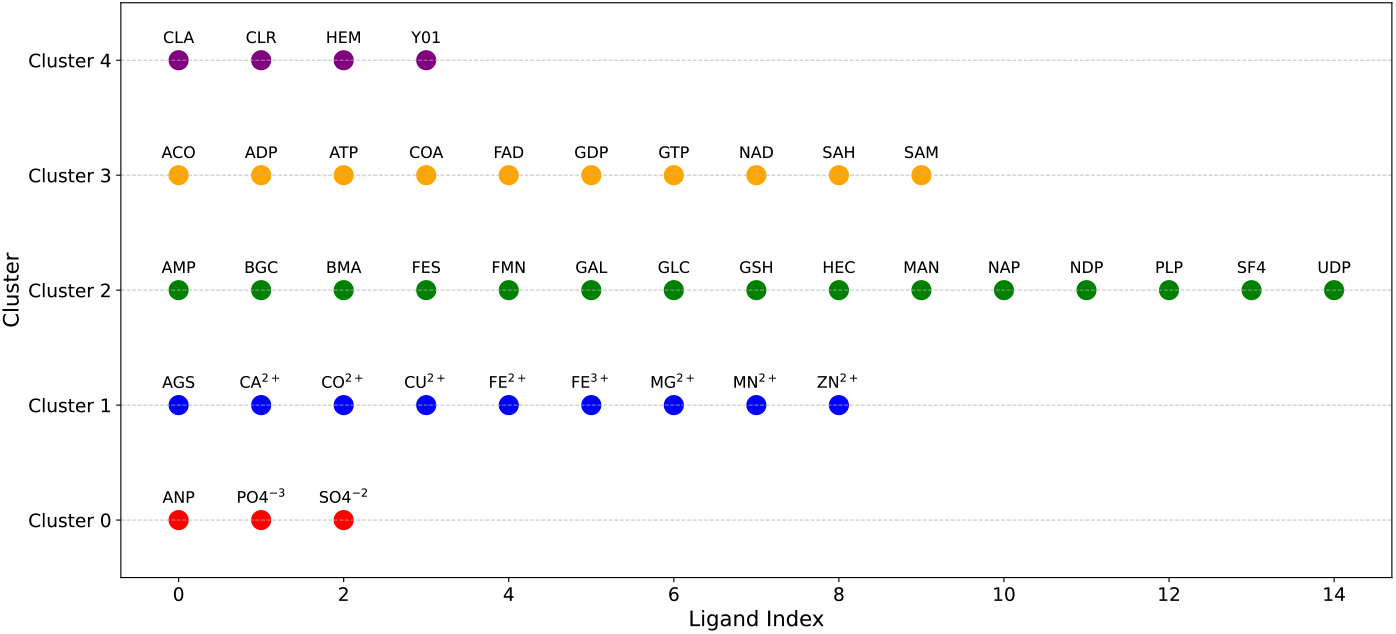
Clustering results of features on the 41 selected ligands.

**Figure 10.**
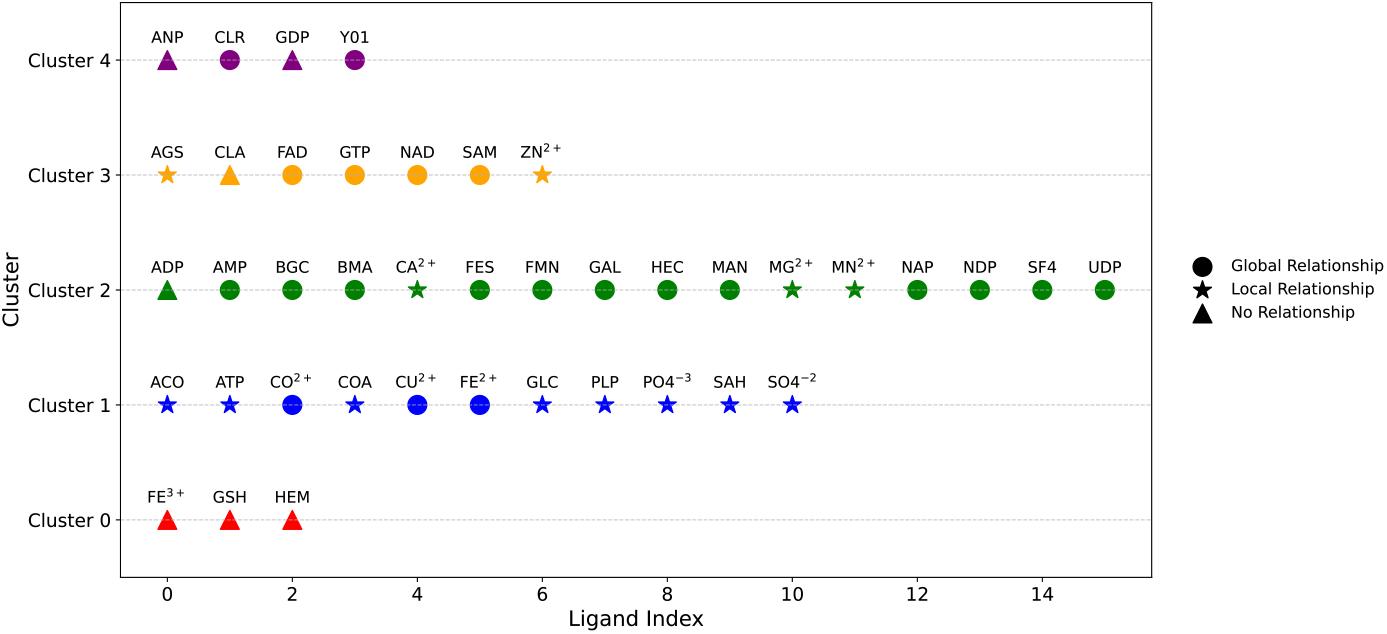
Global Relationships indicate that general biochemical features shared among many ligands have been captured. Local Relationships reflect the successful capture of biochemical properties between specific ligands and their closely related counterparts. No Relationships indicate that the biochemical properties were not captured at all.

**Figure 11.**
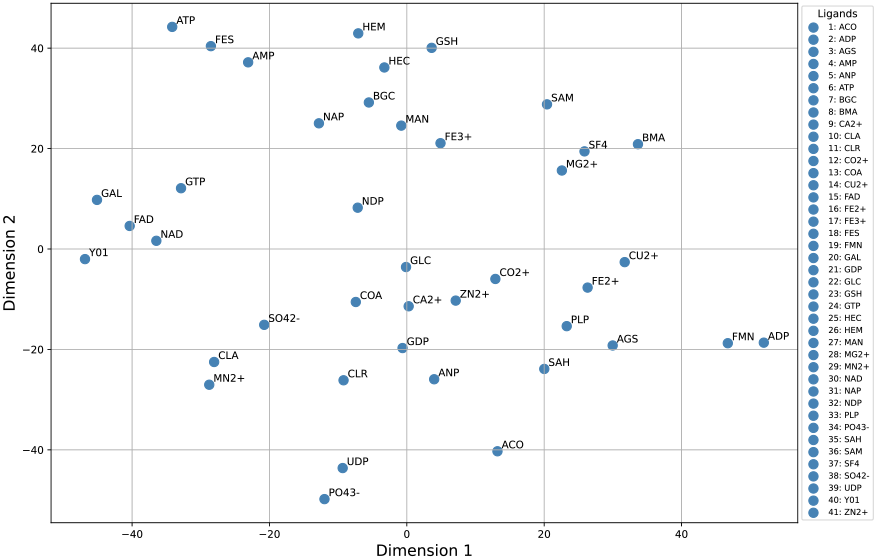
Visualization of task token embeddings using t-SNE.

**Figure 12.**
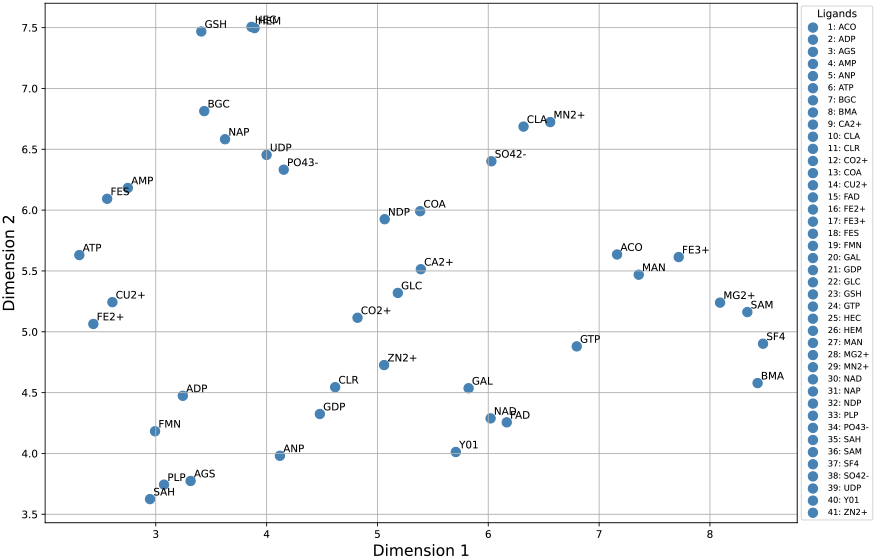
Visualization of task token embeddings using UMAP.

#### A.4.2 Features Pool Creation

The feature pool of 26 descriptors was carefully designed to capture the physical, chemical, and structural properties of ligands, making them particularly suitable for describing protein-ligand interactions. These features were selected using domain knowledge of protein-ligand interactions and their ability to explain binding phenomena effectively. Two primary sources were used to collect these features:

- PubChem, a free online database maintained by the National Center for Biotechnology Information (NCBI), which provides precomputed chemical information for small molecules, drugs, and bioactive compounds. Features were retrieved using Compound IDs (CIDs) and SMILES (Simplified Molecular Input Line Entry System), a text-based representation of molecular structures.
- The second source was RDKit, an open-source cheminformatics toolkit, where SMILES strings were converted into molecular objects and processed using various descriptors to compute additional features.

Table 6 shows the set of 26 features, categorized into seven groups, captures the properties of metal ions and molecules from multiple perspectives, providing a comprehensive description of their binding potential with proteins.

**Table 6:** All 26 features we used in the interpretation step.

| No. | Name | Description | Source |
| --- | --- | --- | --- |
| 1 | MolecularWeight | Total molecular mass | PubChem |
| 2 | ExactMass | High-precision mass of the molecule | PubChem |
| 3 | MolecularVolume | Estimated molecular volume | RDKit |
| 4 | HeavyAtomCount | Count of non-hydrogen atoms | RDKit |
| 5 | RingCount | Total number of rings in the molecule | RDKit |
| 6 | CarbonCount | Number of carbon atoms | RDKit |
| 7 | OxygenCount | Number of oxygen atoms | RDKit |
| **Polarity and Hydrophobicity Features** |  |  |  |
| 8 | LogP | Partition coefficient (hydrophobicity) | PubChem |
| 9 | MolLogP | Alternative measure of hydrophobicity | RDKit |
| 10 | HydrophobicSurfaceArea | Hydrophobic interaction area (TPSA) | RDKit |
| 11 | TPSA | Topological Polar Surface Area (polarity) | PubChem |
| 12 | Hydrophilicity | Difference between molecular weight and hydrophobicity | RDKit |
| 13 | Polarizability | Molecular refractivity | RDKit |
| 14 | Refractivity | A measure of a molecule’s polarizability | RDKit |
| **Charge and Electrostatics Features** |  |  |  |
| 15 | NetCharge | Net electrical charge of the molecule | PubChem |
| 16 | ElectrostaticPotential | Approximate measure of electrostatic potential | RDKit |
| **Flexibility and Rotational Features** |  |  |  |
| 17 | RotatableBonds | Number of rotatable bonds | PubChem |
| 18 | RotatableBondFraction | Fraction of single bonds that are rotatable | RDKit |
| **Bond and Connectivity Features** |  |  |  |
| 19 | SingleBonds | Count of single bonds | RDKit |
| 20 | DoubleBonds | Count of double bonds | RDKit |
| 21 | BalabanJ | Balaban index (topological descriptor) | RDKit |
| **Hydrogen Bonding Features** |  |  |  |
| 22 | HBondDonors | Number of hydrogen bond donors | PubChem |
| 23 | HBondAcceptors | Number of hydrogen bond acceptors | PubChem |
| 24 | HydrogenBondingPotential | Difference between molecular weight and TPSA | RDKit |
| **Aromaticity and - Interactions** |  |  |  |
| 25 | AromaticRings | Number of aromatic rings | RDKit |
| 26 | PiPiInteractionSites | Number of - interaction sites | RDKit |

#### A.4.3 Optimizing Feature Selection

Our approach leverages clustered embeddings as a reference to evaluate clustered features from various feature combinations, identifying the best set of features to describe ligands based on the highest Adjusted Rand Index (ARI) score. We began with task token embeddings of ligands that achieved high F1 scores to ensure noise reduction and high-quality clustering. These embeddings, initially 640-dimensional, were reduced to 27 principal components using PCA while retaining 99% of the variance. The reduced embeddings were then clustered using k-means, with the optimal number of clusters determined via the Elbow method, serving as the target clusters.

To identify the most informative ligand features, we implemented a search algorithm (Algorithm 1) that evaluates all possible combinations of up to 13 features from a pool of 26. In the first iteration, the algorithm selects a single feature (26 possible options). In the second iteration, it selects two features (325 possible combinations). This process continues up to 13 features, yielding approximately 39 million combinations. For each combination, the ligand-based feature clustering is performed, and the ARI score is computed. The feature combination that achieves the highest ARI score is selected as the best set.

Next, we removed the threshold constraint and extended the algorithm to all 41 ligands, examining whether task token embeddings captured meaningful representations for ligands with F1 scores below 0.5. This analysis demonstrated that the embeddings retained significant information even for lower-performing ligands.

Table 7 presents the output of our searching algorithm, showing the top three feature combinations based on the ARI metric for the top 28 ligands. Table 8 displays the top three feature combinations for the entire set of 41 ligands.

**Table 7:** Top three feature combinations for the 28 ligands.

| No. Features | Features | ARI | NMI | Pairwise Accuracy |
| --- | --- | --- | --- | --- |
| 7 | MolecularWeight, NetCharge, RotatableBonds, HydrogenBondingPotential, CarbonCount, SingleBonds, BalabanJ | 0.447 | 0.614 | 0.783 |
| 12 | LogP, NetCharge, RotatableBonds, ExactMass, Polarizability, AromaticRings, MolLogP, MolecularVolume, HydrogenBondingPotential, CarbonCount, SingleBonds, BalabanJ | 0.423 | 0.573 | 0.772 |
| 13 | MolecularWeight, LogP, NetCharge, RotatableBonds, ExactMass, Polarizability, MolLogP, MolecularVolume, RingCount, HydrogenBondingPotential, CarbonCount, BalabanJ, Hydrophilicity | 0.434 | 0.577 | 0.778 |

**Table 8:**
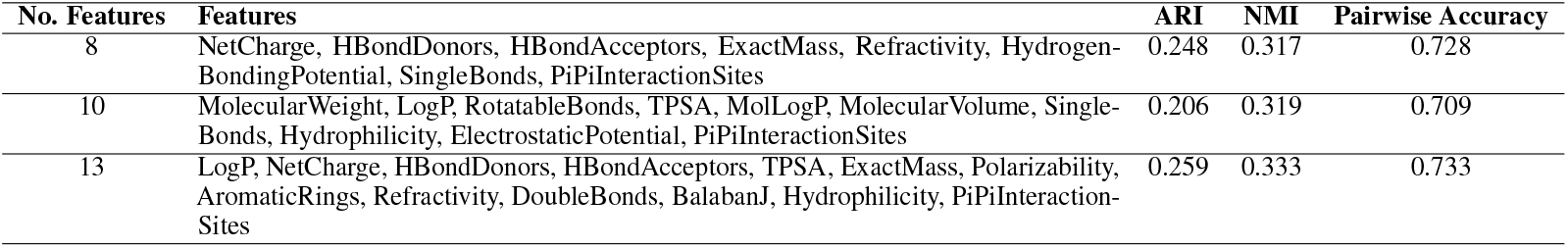
Top three feature combinations for the entire set of 41 ligands.

| No. Features | Features | ARI | NMI | Pairwise Accuracy |
| --- | --- | --- | --- | --- |
| 8 | NetCharge, HBondDonors, HBondAcceptors, ExactMass, Refractivity, HydrogenBondingPotential, SingleBonds, PiPiInteractionSites | 0.248 | 0.317 | 0.728 |
| 10 | MolecularWeight, LogP, RotatableBonds, TPSA, MolLogP, MolecularVolume, SingleBonds, Hydrophilicity, ElectrostaticPotential, PiPiInteractionSites | 0.206 | 0.319 | 0.709 |
| 13 | LogP, NetCharge, HBondDonors, HBondAcceptors, TPSA, ExactMass, Polarizability, AromaticRings, Refractivity, DoubleBonds, BalabanJ, Hydrophilicity, PiPiInteractionSites | 0.259 | 0.333 | 0.733 |

##### Algorithm 1

Ligand interpretation clustering

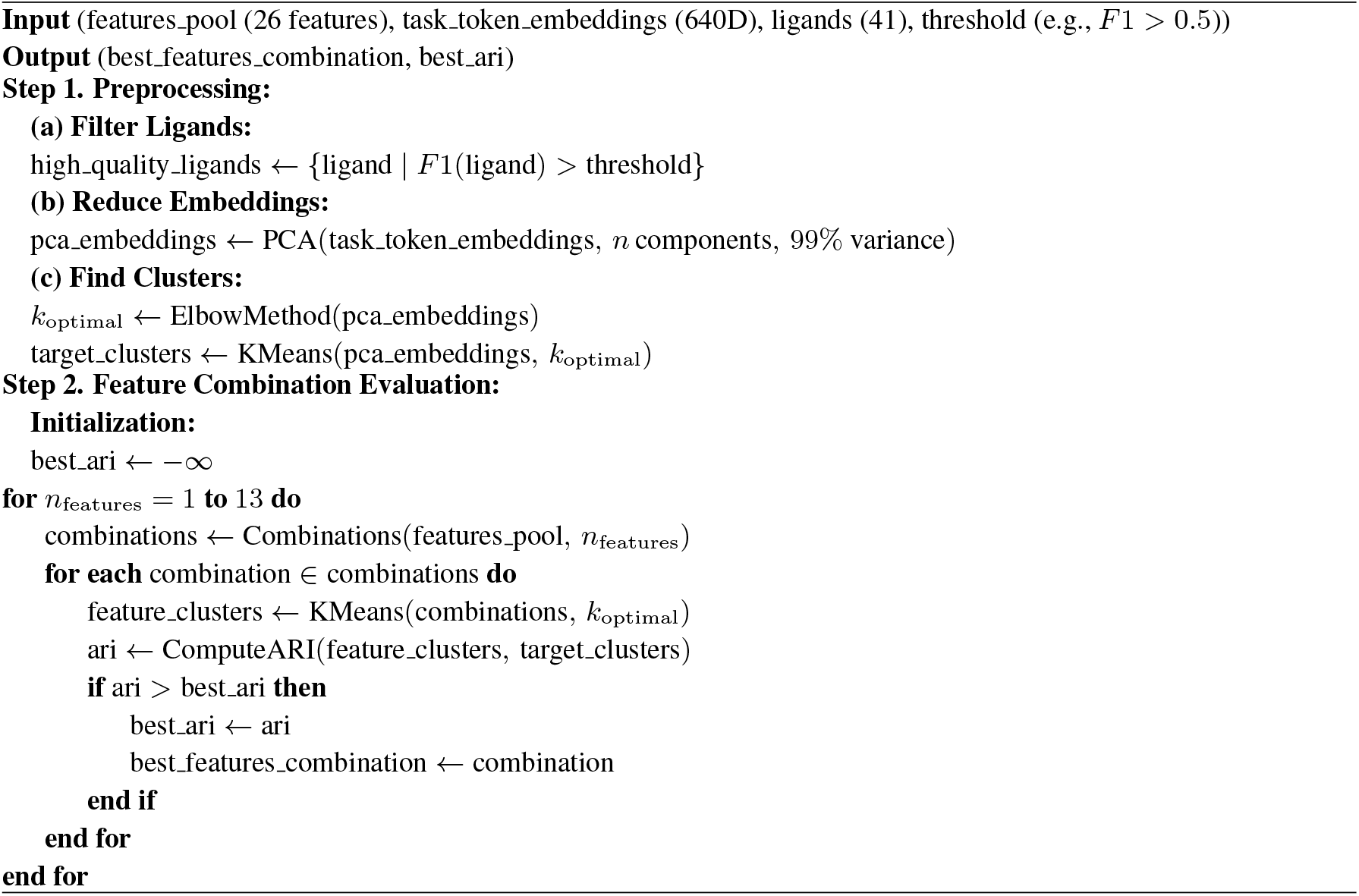

#### A.4.4 Visualization

To analyze the structural relationships within the high-dimensional ligand embeddings, we applied dimensionality reduction techniques to project the representation of 41 ligands from the 640 dimensional into a two-dimensional space for visualization. The methods explored included t-distributed stochastic neighbor encoding (t-SNE) (Figure 11) and uniform manifold approximation and projection (UMAP) (Figure 12).

For the implementation of t-SNE and UMAP, we ensured reproducibility by setting a random state. The perplexity parameter for t-SNE was set to 3, and the number of neighbors (*n neighbors*) for UMAP was also set to 3. These parameters were chosen to focus on capturing local relationships among ligand embeddings and to preserve some global structural details. Additionally, the dimensionality of the output was set to two (*n components* = 2) because the visualizations are in two dimensions. All other parameters were kept at their default settings. Also, we visualized the trained index token embeddings associated with the decoder using t-SNE (Figure 13). We could not see any interpretable patterns in those embeddings.

**Figure 13.**
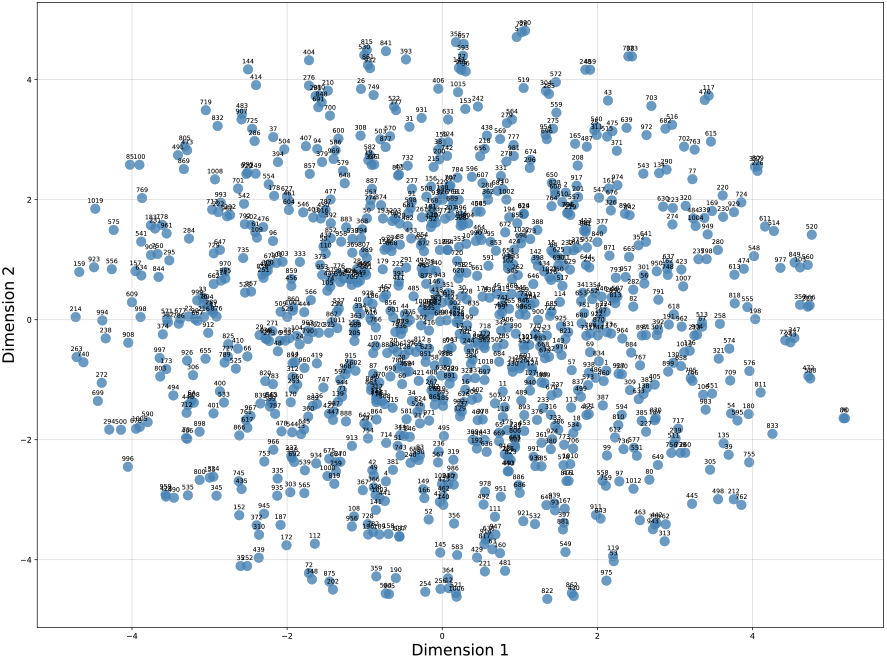
Visualization of index token embeddings using t-SNE.

